# Exercise training in a color-polymorphic lizard reveals differential effects of mating tactics and color morphs on telomere, body condition and growth dynamics

**DOI:** 10.1101/2020.12.23.424255

**Authors:** Christopher R Friesen, Mark Wilson, Nicky Rollings, Joanna Sudyka, Mathieu Giraudeau, Camilla M Whittington, Mats Olsson

## Abstract

Alternative reproductive tactics (ARTs) are correlated suites of sexually selected traits that are likely to impose differential physiological costs on different individuals. While some level of activity might be beneficial, animals living in the wild are often working at the margins of their resources and performance limits. Individuals using ARTs may have divergent capacities for activity, and when pushed beyond their capacity, they may experience condition loss, oxidative stress, and molecular damage that must be repaired with limited resources. We used the Australian painted dragon lizard that exhibits color-polymorphims with corresponding alternative reproductive tactics (ARTs) as a model to experimentally test the effect of exercise on body condition, growth, reactive oxygen species (ROS), and telomere dynamics—a potential marker of stress and aging and a correlate of longevity. For most males, ROS tended to be lower with greater exercise; however, males with yellow throat patches—or bibs— had higher ROS than non-bibbed males. At the highest level of exercise, bibbed males exhibited telomere loss, while non-bibbed males gained telomere length; the opposite pattern was observed in the no-exercise controls. Growth was positively related to food intake but negatively correlated with telomere length at the end of the experiment. Body condition was not related to food intake but was positively correlated with increases in telomere length. These results, along with our previous work, suggest that aggressive bibbed males suffer physiological costs that may reduce longevity.

## Introduction

Many behaviors that are crucial for survival and reproductive success rely on performing stressful, intense, or sustained levels of physical activity, including foraging, escaping from predators, migration, courtship displays, and territorial defense (Arnold, 1983). The link between these critical activities and the investment in life-history and sexually selected traits can involve the energetics of resource allocation (Van Noordwijk and de Jong, 1986), physiological stress (Zera and Harshman, 2001), and trade-offs between self-maintenance, growth, and reproduction (Monaghan, 2014; Pontzer, 2018; Roff, 1992; Soulsbury and Halsey, 2018; Speakman et al., 2015). In humans, it is well established that moderate regular exercise salubriously affects immunity, reproduction, and stress responses (Pontzer, 2018). However, for example, individual humans vary in how they respond to the intensity and duration of exercise training, exhibiting so-called “exercise phenotypes” that are associated with allelic variants at several genomic loci (e.g., Bouchard et al., 2011; Bray et al., 2009; Nickels et al., 2020). An extreme example of how activity influences the expression of life-history, reproductive, and sexually selected traits is hypogonadism, which is common in ultra-endurance and elite athletes (Hackney, 2020; Nickels et al., 2020). It is also generally well established that when humans engage in acute, extremely intense, and prolonged exercise, they suffer from increased oxidative damage (Pontzer, 2018; Powers and Jackson, 2008) and shortened telomeres (Borghini et al., 2015; Denham, 2019; Ludlow et al., 2013; Nickels et al., 2020).

Telomeres are the protective endcaps of chromosomes composed of a repeating sequence of nucleotides—TTAGGG—and associated protein complexes (Blackburn, 1991; Palm and de Lange, 2008). Telomeres shorten with successive cellular divisions *in Vitro* and *in Vivo* (Allsopp et al., 1995; Monaghan and Ozanne, 2018) and are damaged by an excess of reactive molecules (e.g., oxygen-based free radicals, and reactive oxygen and nitrogen species [henceforth ROS for simplicity] (von Zglinicki, 2002). Reactive OS are the natural products of metabolic processes such as the oxidative bursts of immune cells, cellular inflammation responses, and ATP production during oxidative phosphorylation in mitochondria (Reichert and Stier, 2017; von Zglinicki, 2002). ROS-induced damage (oxidative stress), and the consequent telomere erosion, is linked to metabolism, disease, parasite load and inflammation (Asghar et al., 2015; Monaghan, 2010; Monaghan and Haussmann, 2006; Monaghan et al., 2009; Sudyka et al., 2019), physical and psychosocial stress (Bebbington et al., 2017; Haussmann and Heidinger, 2015)), nutrition deficits (Noguera et al., 2015), reproductive effort (Sudyka, 2019; Sudyka et al., 2014), and growth (Geiger et al., 2012; Näslund et al., 2015). The activity of the enzyme telomerase can restore telomere length (Gomes et al., 2010). However, telomerase activity comes at an energetic expense and may also be suppressed in many tissues because telomerase expression can promote oncogenesis (cancer) (Gomes et al., 2010; Olsson et al., 2017).

A key problem in telomere biology that has received less scrutiny is our lack of understanding the factors that underpin individual variation in telomere dynamics and how that might affect or interact with other evolutionary processes. Short telomeres and telomere erosion are candidate biomarkers of the accumulated stressors an individual has endured through life and indicate damage not yet repaired (Monaghan et al., 2018; Wilbourn et al., 2018). These accumulated stressors may link shortening of telomeres with mortality risk (Wilbourn et al., 2018). Indeed, relatively short telomeres are often better predictors of survival than chronological age in some species (Schultner et al., 2014; Tricola et al., 2018). On the other hand, individuals of some species with longer telomeres tend to have higher relative reproductive fitness (e.g., sand lizards, Olsson et al., 2011; Pauliny et al., 2018). All else being equal, there is some evidence that differences in telomere length and shortening among individuals reflects their quality (e.g., high breeding and immune function performance, e.g., Le Vaillant et al., 2015). When individuals vary in character states, trait-associated telomere erosion suggests a correlated cost of those traits and disparate investment in them at the expense of telomere maintenance. For example, painted dragon lizards exhibit a negative correlation between sexually selected colour and telomere maintenance (e.g., Giraudeau et al., 2016; Rollings et al., 2017). Likewise, telomere erosion may be a useful indicator of the cost of intense and sustained activities or behavior (e.g., Sudyka et al., 2014).

Although we may think of wild animals as highly-tuned, high-performance athletes (Irschick and Higham, 2015), their performances incur costs. Fish subjected to only two acute bouts of exhaustive exercise exhibit immediate exercise-induced DNA damage (Aniagu et al., 2006); active and aggression-prone phenotypes may also suffer high telomere erosion (Adriaenssens et al., 2016). In birds, migration reduces antioxidants (Cooper-Mullin and McWilliams, 2016) and shortens telomeres (Bauer et al., 2016; Schultner et al., 2014), and even non-migratory activity increases mortality (Daan et al., 1996; Sudyka et al., 2014). Across vertebrates, endurance and high-intensity activities stimulate the release of glucocorticoids to mobilize energy reserves (e.g., reproductive effort, Bauch et al., 2016). Unless energy reserves are replaced, these activities reduce the availability of those resources for protective and maintenance functions. Glucocorticoids are linked to oxidative stress (Costantini et al., 2011) and to shorter telomeres across different animals (Angelier et al., 2018). Thus, activities that increase evolutionary fitness are likely also to incur costs reflected as telomere erosion.

In lizards, the males of many species patrol and aggressively defend territories (Olsson, 1993; Olsson and Madsen, 1998; Stamps, 1983). Both natural and sexual selection generate directional selection for improved physical performance, such as endurance or sprint speed (Husak and Fox, 2008; Irschick et al., 2008). Faster males are better at defending territories and sire more offspring (Husak et al., 2006; Husak et al., 2008), but performance comes at a cost. Aggressive territorial behavior is energetically expensive and stressful, and it reduces survival (Marler and Moore, 1988; Marler and Moore, 1989; Marler et al., 1995). Resource supplementation mitigates some of these effects, suggesting that resource allocation trade-offs may be an important factor to consider (Marler and Moore, 1991). However, juvenile lizards fed *ad libitum* and experimentally trained for endurance show a reduction in immune function (Husak et al., 2017), and endurance-trained lizards have diminished survival versus sedentary controls when released into the wild (Husak and Lailvaux, 2019).

Color-polymorphic species are a valuable tool in resolving the interplay between sexually selected and life-history traits because the polymorphism is a convenient visual-code for the associated behaviors and physiology of an individual in an otherwise similar genetic background of the species (Stuart-Fox et al., 2020). Here, we test whether the costs of polymorphic mating strategies can be evaluated by experimentally manipulating activity levels and measuring telomere erosion in an Australian color-polymorphic lizard.Male painted dragons (*Ctenophorus pictus*) are polymorphic for head color: red, orange, yellow, and blue (i.e., the same as body color) (Olsson et al., 2007b). The presence or absence of a yellow gular patch, or “bib”, is also a male-polymorphic trait (Olsson et al., 2009a). Maintaining coloration is costly, and color fades over the breeding season, at least partly in response to oxidative stress (Giraudeau et al., 2016; Olsson et al., 2012). These color traits correspond with alternative reproductive tactics and telomere length, which may be mediated by metabolism and oxidative stress (Friesen et al., 2017a; Olsson et al., 2018a; Rollings et al., 2017). In these dragons, both color and oxidative status are at least in part determined by genetics (Olsson et al., 2007b; Olsson et al., 2009d; Olsson et al., 2008). In the lab, mitochondrial superoxide levels (an indicator of oxidative status) are negatively correlated with body condition and endogenous antioxidant activity (Friesen et al., 2019; Friesen et al., 2017b).

To test whether activity level and food consumption may explain morph-specific differences in telomere length and the loss of body condition observed in the wild-caught animals (Healey and Olsson, 2009; Olsson et al., 2009a), we measured telomere erosion in response to manipulated activity (two levels of enforced exercise and a control treatment) over a month. We predicted that the more aggressive morphs (i.e., red-headed males and bibbed males of any head-color) would have higher ROS and suffer more significant telomere erosion than the other morphs (i.e., orange, yellow, and blue males and non-bibbed males of any color) as a result of exercise treatment (Olsson et al., 2018b; Olsson et al., 2017). We predicted that telomere attrition would be most pronounced at the highest level of exercise but would also be mediated negatively by ROS levels and positively by body condition and food consumption. A simple negative relationship between growth and telomere erosion is unlikely given the potentially complex relationships between condition, food intake, and morph-types, and the fact that lizards have indeterminate growth (Shine and Charnov, 1992); nevertheless, cellular replication is often associated with growth and telomere loss (Monaghan and Ozanne, 2018).

## Materials and Methods

### Animal collection

This work was performed under the authority granted by Animal Ethics permit at the University of Sydney (AE20136050), and the lizards were collected under NSW National Parks and Wildlife Service Scientific license (SL100352). Australian painted dragons, *Ctenophorus pictus*, are small (adult snout-vent length 65-95 mm, mass 8-16 g) diurnal lizards of sandy habitats and low vegetation, with a range covering central New South Wales to Western Australia (Cogger, 2014). *Ctenophorus pictus* are annuals, with nearly 90% dying within one year in this population (Olsson et al., 2007b). We caught 56 mature (~ 9 months old) male lizards by noose or hand at Yathong Nature Reserve, NSW, Australia (145°35’E; 32°35’S) during the Australian spring (ca. mid-October, early in the breeding season). We exhaustively sampled with daily repeated passes along a ~ 15 km transect over seven days; hence, our sample should approximately represent natural head color and bib-morph frequencies in the wild at the time. Head color (HC): red n = 15 (26.8%); orange n = 13 (23.2%); yellow n = 17 (30.4%); blue n = 11 (19.6%). Bib-morphs: bibbed n =20 (35.7%); non-bibbed n =36 (62.3%). The animals were held in facilities at the University of Sydney where they were housed individually in opaque plastic enclosures (330×520×360 mm) with a sandy substrate and exposed to a 12:12 h light:dark light-cycle. We fed each lizard exactly four mealworms (of the same size class) lightly dusted with calcium and multivitamins, every other day. We misted the enclosures with water once per day. Before each feeding, we removed and tallied the number and proportion of uneaten worms, which allowed us to calculate food consumption throughout the experiment. Heat lamps and ceramic hides were provided to allow the lizards to thermoregulate to their preferred body temperature (~35-36 °C (Melville and Schulte II, 2001)).

### Exercise treatments

Head-color and Bib-morphs were distributed among one of three endurance exercise treatment groups: No exercise (0X; N = 18); light exercise (1X, N = 19); and heavy exercise (3X, N = 19). Endurance exercise was similar to a previous protocol (Tobler et al., 2012; Uller and Olsson, 2003; Uller and Olsson, 2006), which consisted of letting the lizards swim in a water bath (60 cm × 60 cm and 20 cm deep) of constant temperature (34-36 °C, monitored using a thermometer). Lizards were individually transferred directly from their home enclosures into the water, where they naturally began to swim using an undulatory motion. When a lizard stopped swimming, he was persuaded to continue with a light tap to the tail. If, after three consecutive taps, the lizard did not resume swimming, he was considered to be exhausted and returned to his home enclosure. Light exercise males (1X) swam once to exhaustion before removal to the home enclosure. Heavy exercise males (3X) swam to exhaustion three times with 30 minute recovery periods spent in their home enclosure between swims. No exercise males (0X) were used to control for handling effects: each was caught, picked up, and placed in a water bath (60 × 60 cm and 2 cm deep) of water (34-36 °C) water for 3 minutes (~average of an exhaustive swim), and returned to their cage. All treatments (0X, 1X, 3X) were administered three days per week for approximately four weeks, beginning 4 November and ending 3 December. This period corresponds to the mid-to-late period of the breeding season.

To ensure that no exercise treatment group had an overabundance of “high-endurance” exercise phenotype lizards, we timed all males during exercise to exhaustion the day before allocating them to a group (there was no difference pre-allocation between treatment groups in the time until exhaustion, F_2,53_ = 1.803, P = 0.175). We collected blood samples and recorded body mass (to 0.01 g) and snout-vent length (SVL; to 1 mm) before the experiment commenced (late October) and after the experiment was completed (early December). The blood samples were used for telomere and ROS measurements. The same experienced operator took the SVL measures. Three males died (one from 3X and two from 0X) at different points in the experimental period (no significant effect of treatment), leaving 53 survivors. This level of mortality (~ 5%) is typical in the lab for this short-lived dragon (Friesen and Olsson pers. obs.).

### Sample collection

#### Blood sampling

We sampled blood (~ 100 μL) from each lizard before and after the completion of the treatment period by using a capillary tube and gently perforating the *vena angularis* (in the corner of the mouth). An aliquot of peripheral blood (10 μl) from each male was diluted with 9 volumes of phosphate-buffered saline (PBS; 137 mM NaCl, 2.7 mM KCl, 1.5 mM KH_2_PO_4_, 8 mM Na_2_HPO_4_, pH 7.4) and stored on ice prior to ROS analysis (see below). The remaining blood was centrifuged, the plasma removed, and the remaining pellet of nucleated red blood cells was resuspended in 200 μL of PBS and stored immediately at -80°C for later use in qPCR quantification of telomeres (we avoided the top layer or “buffy-coat” of the pellet that sits above the red blood cells, as it contains the white blood cells and may affect telomere length measurement (Olsson et al., 2020)).

### Quantifying relative telomere length (rTL)

These methods have been described in detail elsewhere (Rollings et al., 2017; Rollings et al., 2014). In brief, we purified DNA from 50 μL blood using a DNeasy Blood and Tissue Kit (Qiagen, Australia), according to the manufacturer’s instructions. RNase A (Qiagen, Australia) was added at the recommended concentration. The DNA concentration (ng/μL) of each sample was measured in duplicate using a Pherastar FS (BMG, Labtech, Germany) and aliquots diluted to 10 ng/μL using the AE buffer provided in the DNA extraction kit. Telomere length was measured using real-time quantitative PCR (qPCR) and a SensiMix SYBR No-ROX Kit (Bioline, Sydney, Australia). The telomere primers used were Tel1b (5’-CGGTTTGTTTGGGTTTGGGTTT GGGTTTGGGTTTGGGTT-3’) and Tel2b (5’-GGCTTGCCTTACCCTTACCCTTACCCTTACCCTTACCCT-3’) (Criscuolo et al., 2009). The 18S ribosomal RNA (*18S*) gene (92 bp amplicon in *Anolis*) was selected as the reference gene, as it had previously been validated in a reptile (Plot et al., 2012) including this species (Rollings et al., 2017). The primer sequences used were 18S-F (5’-GAGGTGAAATTCTTGGACCGG-3’) and 18S-R (5’-CGAACCTCCGACTTTCGTTCT-3’). The melt curves produced for both telomere and 18S after amplification by qPCR displayed a single peak, indicating specific amplification of the target DNA sequence. The qPCR was performed in a final volume of 20 μl for both telomeres and 18S. Ten ng of DNA was used per reaction, and the primers were used at a concentration of 250 nM. For each reaction, 11.25 μl SensiMix SYBR No-ROX Master Mix (Bioline, Australia) was added, and MgCl_2_ was added for a reaction concentration of 1.7 mM. Reactions were run in triplicate for each sample. Amplifications were carried out in a Rotor-Gene 6000 thermocycler (Qiagen, Australia) using an initial Taq activation step at 95 °C for 10 min, and a total of 40 cycles of 95 °C for 15 s, 60 °C for 15 s and 72 °C for 15 s. A melt curve was created over the temperature range of 60 to 95 °C after each run to ensure no non-specific product amplification. No-template control reactions were run in triplicate for each primer set during every qPCR run to ensure no contamination. Standard curves were created using the blood of a randomly selected lizard, for both telomeres and 18S, to ensure consistent rates of amplification over a wide range of DNA concentrations. The reaction was considered consistent when a straight line with an *R^2^* exceeding 0.985 could be fitted to the values obtained. The efficiency of the telomere amplification was 1.05, and the efficiency of the 18S amplification was 0.96. LinRegPCR 2015.2 (Ruijter et al., 2009a; Ruijter et al., 2009b; Ruijter et al., 2009c; Tuomi et al., 2010) was used to analyze the qPCR data.

### Quantifying superoxide and other reactive oxygen species (ROS)

Mitochondrial superoxide and general reactive oxygen species analyses were completed within 4 h of sampling blood. Before staining, diluted blood was diluted a further 50-fold with PBS and then centrifuged (300 g for 5 min) to pellet cells; each cell pellet corresponded to 10 μl of whole blood. Cells were resuspended in 100 μl of PBS containing either no additions (unstained control), 5 μM MitoSOX Red (MR; Thermofisher; detects superoxide), or 0.1 mM dihydrorhodamine 123 (DHR; Thermofisher; detects reactive intermediates such as peroxide and peroxynitrite).

The fluorescence from these probes can be used to detect, respectively, superoxide ions (MR) and hydrogen peroxide and peroxynitrite (DHR). MR and DHR were added from stock solutions in dimethylsulfoxide (DMSO); the final concentration of DMSO was 0.2 % (v/v) or less. Cells were subsequently incubated at 36°C for 30 min, then washed with PBS by centrifugation as described above and held on ice until analysis by flow cytometry. A total of 50,000 events were acquired for all samples. Flow cytometry was performed using a Becton Dickinson LSRFortessa X20, with excitation at 488 nm for both MR and DHR, and emitted fluorescence was collected using bandpass filters of 575 +/− 13 nm (MR), 515 +/-10 nm (DHR). Data were acquired and analyzed using FACSDiva v4.0.1 (Becton Dickinson, Sydney, Australia) and FloJo (v9.1; TreeStar Inc., USA) software, respectively. On the basis of forward angle laser scatter and side angle laser scatter, a number of blood cell populations were discerned; the results obtained were similar for all these populations. For each sample, the arithmetic mean fluorescence for all 50,000 cells acquired was determined using FloJo software and used to compare between samples and treatments. The accuracy and consistency within a sample period of flow cytometry results have been validated in our previous work (r = 0.97, (P < 0.001), see Olsson et al., 2008for further details).

### Data preparation and statistical analyses

Body condition (BCI) was calculated as the residuals from a regression analysis of mass as a function of SVL, which renders BCI independent of body length (*r* = -0.138, P = 0.325). Body condition is useful because it positively correlates with energy reserves and the endogenous antioxidant superoxide dismutase (SOD) (Friesen et al., 2019; Friesen et al., 2017b). Superoxide measurements were collected on successive days within both sampling periods (late Oct and early Dec); thus, we standardized measurements (z-transformation) within the day the measurements were taken to account for potential batch effects (Friesen et al., 2019; Nakagawa et al., 2017).

The dependent variables, relative telomere length (rTL), body condition (BCI), body length (SVL), body mass (Mass), superoxide (SOx), general ROS (ROS, hereafter), were tested for the effects of exercise on the change from the beginning to the end of the experimental period (∆ = final/end – initial/beginning value), e.g., ∆rTL, ∆BCI, and ∆SVL values, but we also tested for correlations among the ∆ variables, like ∆rTL vs. ∆BCI, as well as correlations like ∆rTL vs. food intake and superoxide and general ROS.

We first tested for correlations between variables to explore how variables might interact to explain changes in ∆rTL. We used both Kendall’s tau and Spearman’s rho non-parametric tests, (using their congruence as a test of sensitivity) when either variable in a correlation test failed normality as determined by Kolmogorov-Smirnov tests. Both Spearman’s rho and Kendall’s tau correlations are reported when they give different results in terms of significance at α ≤ 0.05. Otherwise, Pearson correlations are reported when both variables passed tests of normality. We also used non-parametric tests to explore differences in the dependent variables between morphs (Kruskal-Wallis (k number of groups >2) and Mann-Whitney (k=2)) and exercise regimes (Jonckheere-Terpstra post-hoc test for ordered differences in treatment effects).

We conducted GLM analyses to investigate the effect of exercise treatment on ∆rTL, ∆BCI, and ∆SVL, ∆ROS, and ∆ superoxide (∆SOx) as well as these variables’ associations with head-color and bib-morphs and morph x exercise interactions. We did not include a three-way interaction because of sample size. We constructed initial “full” GLM models with exercise, head-color, bib, exercise x head-color, and exercise x bib as fixed factors. To avoid drawing inferences from an over-fit model, variables with the highest Type III SS P-values were stepwise, backward eliminated from the fullest initial model until only variables with P < 0.2 remained. Then we further reduced the model by only retaining variables that improved model fit as indicated by a ∆AICc (Akaike information criterion for small sample size) of -2.0 or better, stopping when the model was not improved by removing variables (Konishi and Kitagawa, 2008). If an interaction was retained, then we also included both fixed terms of the interaction regardless of their statistical significance. Pairwise posthoc analyses employed the Benjamini-Hochberg procedure to control false discovery rates, but the full tables are included as supplementary materials. In figures, we present standardized (mean = 0, SD = 1) GLM-estimated means and 95% confidence intervals as error bars (Nakagawa et al., 2017; Verhulst, 2020). We also include a dashed line on the z-score scale that indicates “0”, or no change, on the original scale.

With the exception of ∆SVL, all of the “∆” response values adhered to assumptions of normality (Kolmogorov-Smirnov tests, all P > 0.200: ∆SVL K-S Dist. = 0.196, P < 0.001). ∆SVL was further evaluated by visual inspection (histogram Q-Q and P-P plots) and by generating the kurtosis (1.6197) and skewedness statistics (0.7833, SE = 0.3274:). Positive kurtosis greater than zero indicated that the ∆SVL data exhibited more extreme outliers than a normal distribution. This distribution of ∆SVL had significant (skewedness > 2 x SE) positive skewness has a long right tail. Q-Q plots generated by fitting ∆SVL to a gamma distribution indicated a good fit, so we modeled ∆SVL with a GLM (gamma distribution, log link function) after adding 0.2mm to each ∆SVL measurement to eliminate zeros from the dataset to conform with the assumptions of a gamma distribution GLM, but we plot raw values including “negative growth” in SVL. We note that while measurement error may in part explain why we have “negative growth” in ~ 20% of the males, however, there is precedent for shrinkage in lizards (Wikelski and Thom, 2000), the same experienced person measured all lizards, and measurement errors would be randomly distributed across telomere lengths (and morphs). We report the median, range, and % change of initial SVL and the GLM estimated means and 95% confidence intervals in original units.

All statistical analyses were performed in SPSS 25.0 (IBM, Armonk, NY, USA) and Sigma Plot 13.0 (Systat Software Inc. San Jose, CA, USA).

## Results

### Morph-specific change in relative telomere length (∆rTL)

Relative telomere length was rank-consistent between an individual’s measurement before and after exercise treatment (Spearman’s ρ = 0.714, P < 0.001; ICC = 0.880, 0.800-0.929 95%CI, P < 0.001).

The patterns of ∆rTL depended significantly on the interaction between exercise treatment and each morph type (the full model better fit the data than the exercise treatment-only model ∆AICc = - 30.58: **Table 1**). Non-bibbed males exhibited telomere erosion at the 0X level of exercise but gained telomere length at the 1X and 3X levels of exercise, and bibbed males exhibited the opposite relationship (**Fig 1A**; S1 pairwise comparison tables). The interaction of exercise and head-color was not straightforward, so we give a qualitative description. Among red males, ∆rTL was not significantly different across the levels of exercise. Blue males tended to gain telomere length in the no exercise treatment (0X), but lost or maintained telomere length with exercise (1X and 3X). Orange males lost telomere length at the 0X and 3X levels but gained telomere length slightly at the 1X level of exercise. Yellow males lost or maintained telomere length at the 0X and 1X levels but gained slightly under the 3X exercise treatment (**Fig 1A;**S1 tables)).

**Table 1:** ∆ rTL. Results of GLM (normal, identity link function). Bold text in p-value column indicates significance at P ≤ 0.05.

| *Source* | **Likelihood** |  |  |
| --- | --- | --- | --- |
|  | **ratio $X^2$** | **df** | **P-value** |
| ***Full model vs intercept only model*** | 25.356 | 14,38 | **0.031** |
| *Tests of model effects* |  |  |  |
|  | **<u>Wald $X^2$</u>** | **<u>df</u>** | **<u>Type III</u>** |
| *Exercise Treatment* | 1.341 | 2,38 | .512 |
| *Head-color morph* | 5.408 | 3,38 | .144 |
| *Bib* | .059 | 1,38 | .808 |
| *Exerc.Treat. \* HC-Morph* | 47.532 | 6,38 | **<0.001** |
| *Exerc.Treat. \* Bib* | 19.330 | 2,38 | **<0.001** |

**Figure 1.**
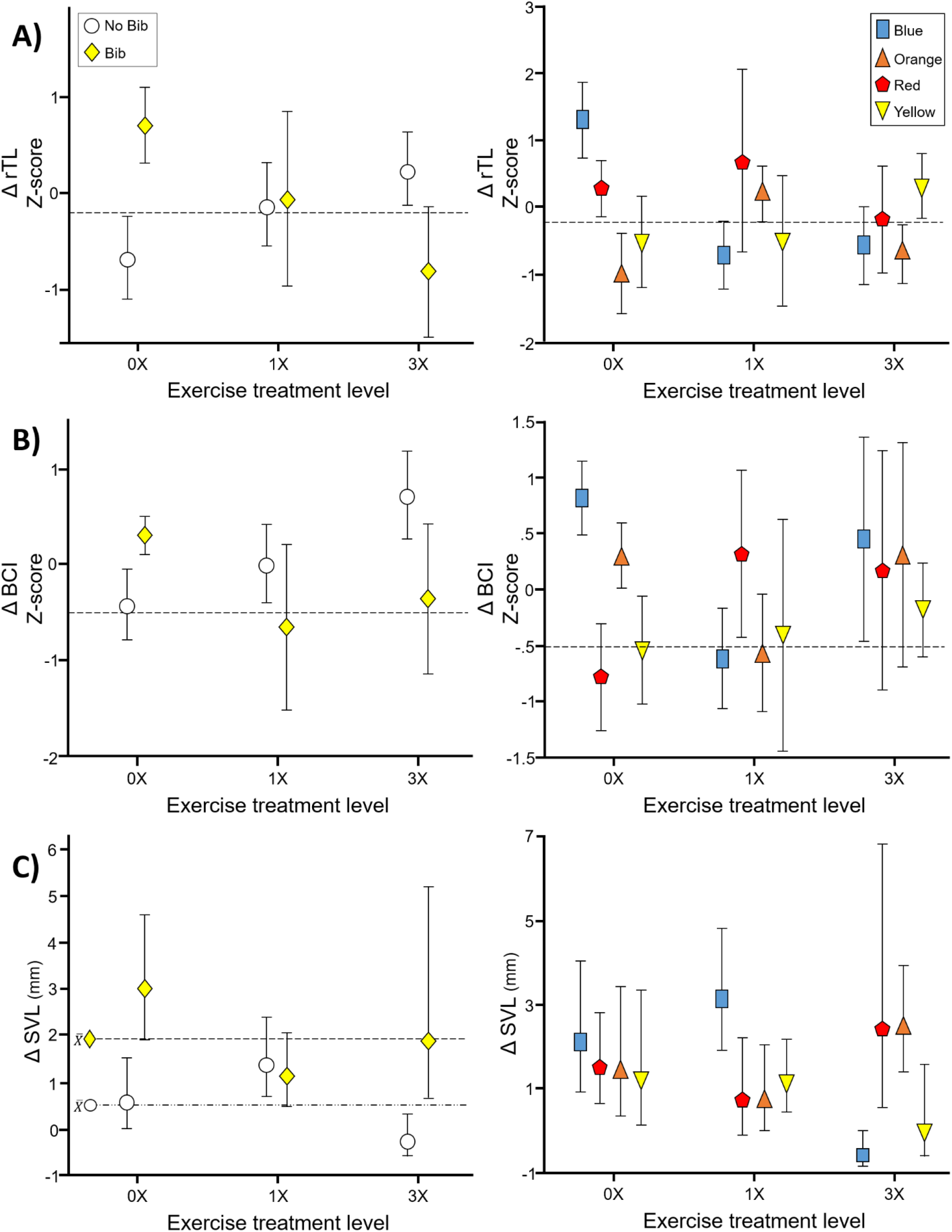
Interaction plots of exercise and morph-types of painted dragons: change in A) telomere length (∆ rTL), B) body condition (∆ BCI), and C) growth (∆SVL). Plots on the left are of bib-morphs, and those on the right are head-color morphs. ∆ rTL and ∆ BCI have been standardized (z-transformation), and ∆ SVL is plotted as the raw untransformed difference in SVL (mm). All symbols are centered on the estimated mean, and the bars represent 95% confidence estimates of the mean from generalized linear models. The dashed horizontal lines in plots A and B represent “0” or “no change” on the original unstandardized scale. The dashed horizontal lines in C, for bib-morphs, represents the overall mean of each morph as indicated by the *X̅* next to the symbol.

### Condition and growth

#### Food consumption

Males ate most of the food offered to them (median = 91.1% excluding one male that ate only 26.7% of his food. The food eaten by the remainder of the males ranged from 66.7 to 100%). There was no difference in the percentage of food consumed across exercise treatments (Jonckheere-Terpstra test: z df3 = -1.487, P = 0.137). There was weak evidence of a difference among morphs (Kruskal-Wallis H = 7.752, P = 0.051, but group medians ranged: 86.7-97.8%). There was no evidence of a difference between bib-morphs (Mann-Whiteny U = 346.5, P = 0.661). Food consumption was not significantly correlated with ∆rTL (Spearman’s ρ = -0.214, P = 0.124) or ∆BCI (Spearman’s ρ = 0.046, P = 0.741) but was positively correlated with ∆SVL (Spearman’s ρ = 0.295, P = 0.032). There was mixed evidence that ∆SOx (see below) and ∆ROS were positively correlated with food consumption depending on the statistical test used (∆SOx, Spearman’s ρ = 0.262, P = 0.058; Kendall’s τ = 0.201, P = 0.041; ∆ROS, Spearman’s ρ = 0.263, P = 0.058).

#### Body condition change (∆BCI)

Body condition index rank was consistent between an individual’s measurement before and after exercise treatment (Spearman’s ρ = 0.723, P < 0.001; ICC = 0.661, 0.414-0.806 95%CI, P < 0.001).

Overall, across all males, body condition tended to increase through the treatment period (paired t-test; t52 = 3.705, P < 0.001); however, visual inspection of the data indicated some males lost while others gained condition. ∆BCI was positively correlated to ∆rTL (r = 0.316, P = 0.021, **Fig 2**).

**Figure 2.**
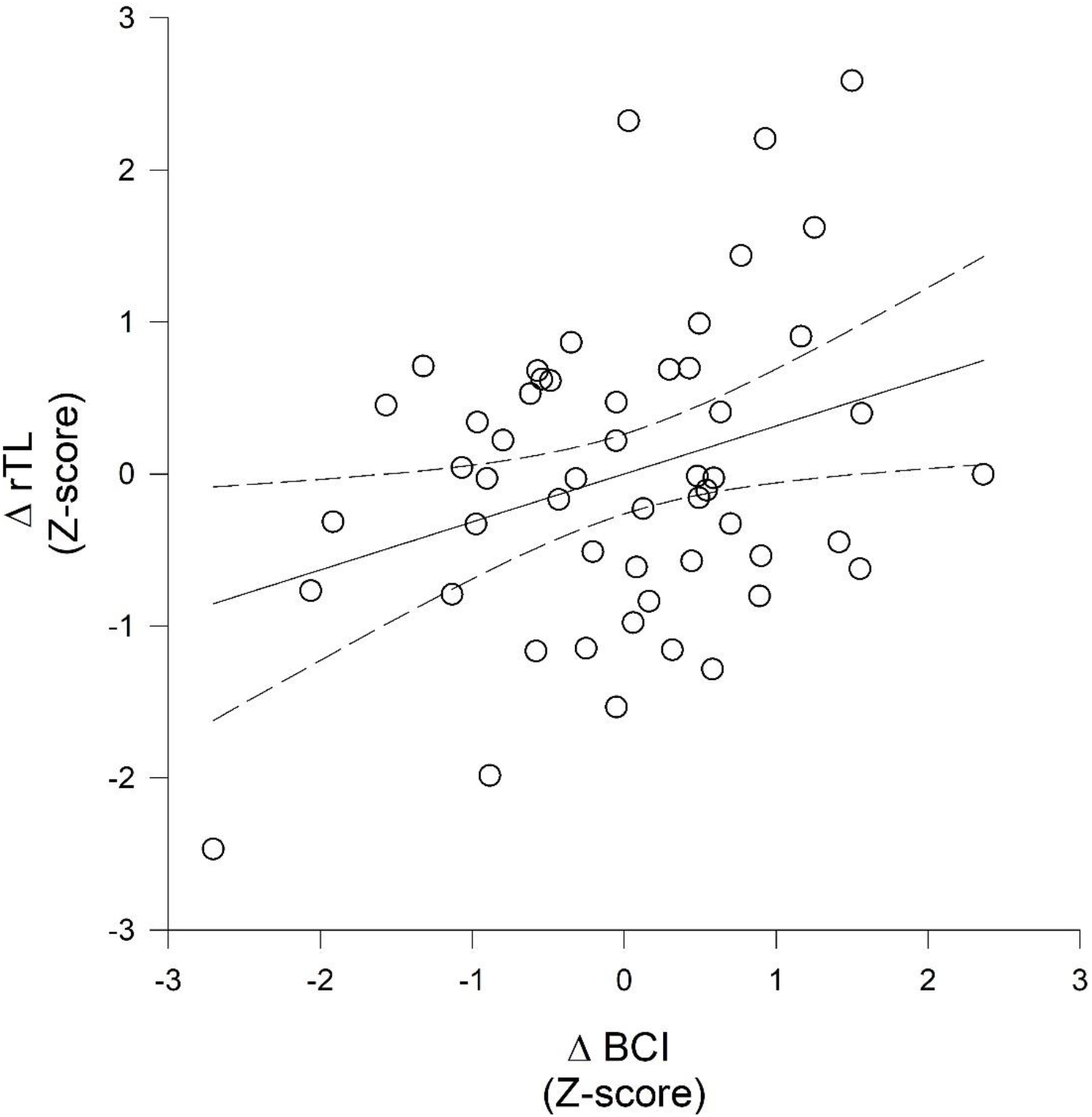
Relationship between the change in relative telomere length (∆rTL) and change body condition (∆BCI) of painted dragons from the beginning to end of the exercise period (r = 0.316, F_1, 51_ = 5.654, P = 0.021). Both variables are standardized (z-transformation).

∆BCI was also affected by exercise treatment through interactions with head-color and bib-morph (**Table 2**). The patterns of ∆BCI across exercise treatments in both bibbed and non-bibbed were congruent with the patterns in ∆rTL. That is, for bibbed males, the trend in ∆BCI 0X > 1X >3X but 0X < 1X < 3X in non-bibbed males (**Fig 1B**). Similarly, the patterns of ∆BCI and ∆rTL were congruent across exercise treatments in blue morphs, but there was no clear pattern of ∆BCI across exercise treatments in the other three morphs (**Fig 1B**; S1 tables).

**Table 2:**
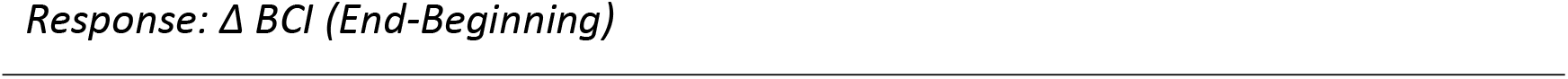
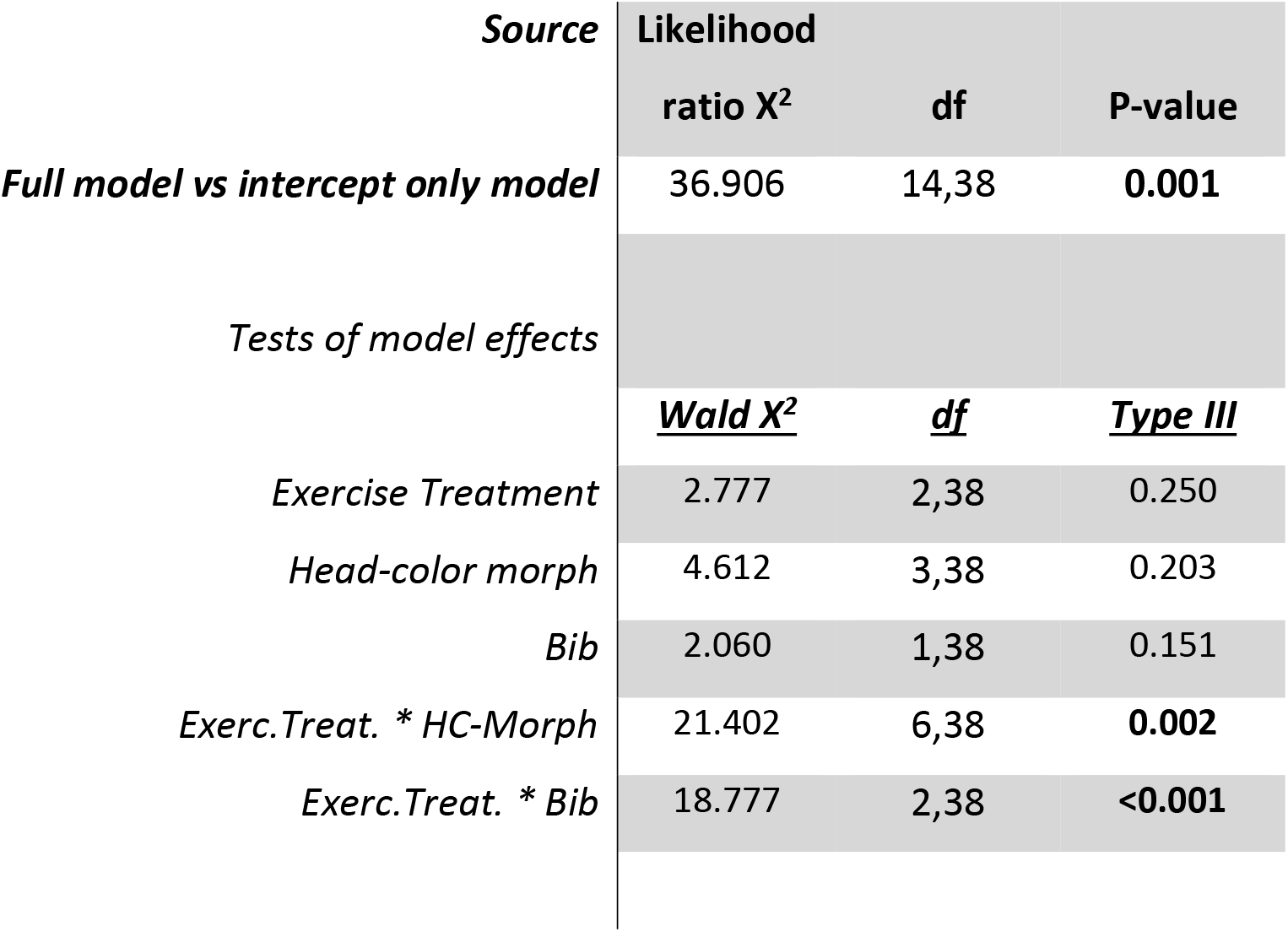
∆ BCI. Results of GLM (normal, identity link function). Bold text in the p-value column indicates significance at P ≤ 0.05.

| *Source* | **Likelihood** |  |  |
| --- | --- | --- | --- |
|  | **ratio X<sup>2</sup>** | **df** | **P-value** |
| ***Full model vs intercept only model*** | 36.906 | 14,38 | **0.001** |
| *Tests of model effects* |  |  |  |
|  | **<u>Wald X<sup>2</sup></u>** | **<u>df</u>** | **<u>Type III</u>** |
| *Exercise Treatment* | 2.777 | 2,38 | 0.250 |
| *Head-color morph* | 4.612 | 3,38 | 0.203 |
| *Bib* | 2.060 | 1,38 | 0.151 |
| *Exerc.Treat. \* HC-Morph* | 21.402 | 6,38 | **0.002** |
| *Exerc.Treat. \* Bib* | 18.777 | 2,38 | **<0.001** |

#### Growth (∆SVL and ∆Mass)

Growth in length (∆SVL) was not correlated with ∆BCI (Spearman’s ρ = -0.171, P = 0.222) nor with change in body mass (∆Mass: Spearman’s ρ = -0.091, P = 0.516). The ∆SVL and ∆rTL were not correlated (Spearman’s ρ = 0.010, P = 0.975); however, ∆SVL was negatively correlated with rTL at the *beginning* of the experiment (initial rTL, Spearman’s ρ = -0.341, P = 0.012) and with rTL at the end of the exercise treatment period (final rTL, Spearman’s ρ = -0.278, P = 0.044: **Fig 3)**.

**Figure 3.**
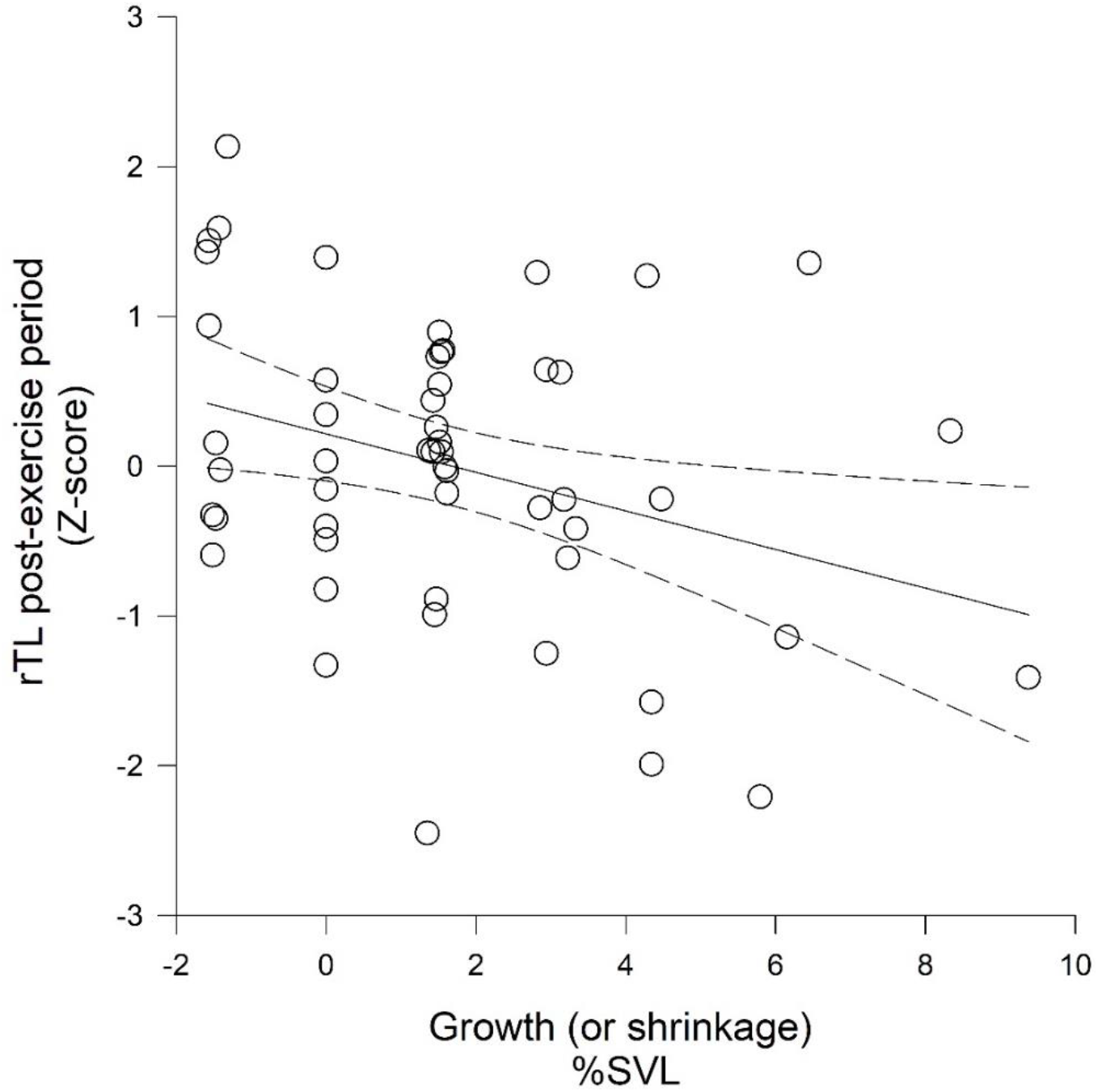
The relationship between relative telomere length (rTL) measured at the end of the exercise period (z-transformed) and the change in snout-to-vent (SVL) length expressed as a percentage of initial SVL in painted dragons. Transforming raw change in SVL (∆mm) to a percentage of the initial SVL improved normality of the residuals, satisfying assumptions of linear regression (K-S test, P = 0.777, r = 0.326, F_1,51_ = 6.079, P = 0.017).

Across males, 10 individuals were 1 mm *shorter*, nine did not grow in length, but the majority (N = 34) gained SVL through the treatment period (Wilcoxon signed-rank test, SVL_2_-SVL_1_: W = 746, Z = 4.396, P < 0.001); the distribution was right-skewed towards higher values (median ∆SVL = 1 mm, range of 7 mm (-1 - 6mm; -1.59 - 9.38% of initial length). Overall, bibbed males exhibited greater growth in length than non-bibbed males (Mann-Whitney rank-sum test, ∆SVL: U = 198, T = 638, P = 0.018). Eight of the 10 males that shrunk in length were non-bibbed males (HC morphs: 3 blue; 3 red; 0 orange; 4 yellow). In addition, 10 males (HC morphs: 3 blue; 1 red; 3 orange; 3 yellow) had ≥ 4% increases in SVL, and 8 of these males had bibs.

∆SVL was also affected by exercise treatment alone (post hoc Jonckheere-Terpstra test: z _df 3_ = -2.121, P = 0.034; ∆SVL, 0X = 1X < 3X) and through interactions with head-color and bib-morphs (**Table 3**). The pattern of ∆SVL across exercise treatments in bibbed males was qualitatively similar to the patterns of ∆rTL. Bibbed males grew the most in the 0X treatment, in which rTL also increased, but grew much less in the 1X and 3X than the 0X treatment. Non-bibbed males grew the least, with some males losing length in the 3X treatment, in which they gained the most in rTL. ∆SVL and ∆rTL exhibited qualitatively opposite patterns across exercise treatments in blue morphs, but no clear pattern for ∆SVL was apparent in the other three morphs (**Fig 1C)** (S1 tables).

**Table 3:** ∆ SVL. Results of GLM (gamma, log link function). Deviance value = 36.639, degrees of freedom = 38, Deviance/df = 0.964, indicating that the model is not over-dispersed. Bold text in the p-value column indicates significance at P ≤ 0.05.

| *Source* | **Likelihood** |  |  |
| --- | --- | --- | --- |
|  | **ratio X<sup>2</sup>** | **df** | **P-value** |
| ***Full model vs intercept only model*** | 25.212 | 14,38 | **0.033** |
| *Tests of model effects* |  |  |  |
|  | **<u>Wald X<sup>2</sup></u>** | **<u>df</u>** | **<u>Type III</u>** |
| *Exercise Treatment* | 5.605 | 2,38 | 0.061 |
| *Head-color morph* | 3.489 | 3,38 | 0.322 |
| *Bib* | 10.103 | 1,38 | **0.001** |
| *Exerc.Treat. \* HC-Morph* | 27.388 | 6,38 | **<0.001** |
| *Exerc.Treat. \* Bib* | 9.658 | 2,38 | **0.008** |

Across all males, body mass did not systematically change through exercise treatment period (paired t-test; t_52_ = 1.115, P = 0.270); likewise, neither exercise treatment, morph-type, nor their interaction were associated with ∆Mass (all P > 0.087) nor was ∆rTL associated with ∆Mass (r = -0.130, P = 0.353).

#### Reactive oxygen species: Superoxide and other ROS

##### Superoxide

Superoxide was not consistent between an individual’s measurement before and after exercise treatment (ICC = -0.069, -0.331-0.202 95%CI, P = 0.691), and change in superoxide (∆SOx) through the exercise treatment period was not related to head-color or bib-morph or their interaction with exercise (all P > 0.109). However, superoxide at the end of exercise treatment (*final* SOx) depended on exercise treatment, head-color, and bib-morph (**Table 4; Fig 4**) but not the interactions between exercise treatment and either of the morph types (backward eliminated, P > 0.230). Bibbed males had higher *final* SOx than non-bibbed males (Bib: P = 0.038); *final* SOx was significantly lower in the highest exercise (3X) group than in the no exercise (0X) group (Exercise: P = 0.049); and orange males had lower *final* SOx than all other morphs (HC-morph, P = 0.032; all pairwise P < 0.048), which was the only difference among head-color morphs. *Final* SOx was weakly, non-significantly, negatively related to ∆BCI (r = - 0.239, P = 0.085) and positively with growth (∆SVL, Spearman’s ρ = 0.250, P = 0.071).There was no evidence that *final* SOx was directly related to relative telomere length (∆rTL: r = 0.049, P = 0.607; final rTL: r = -0.056, P = 0.555).

**Table 4:**
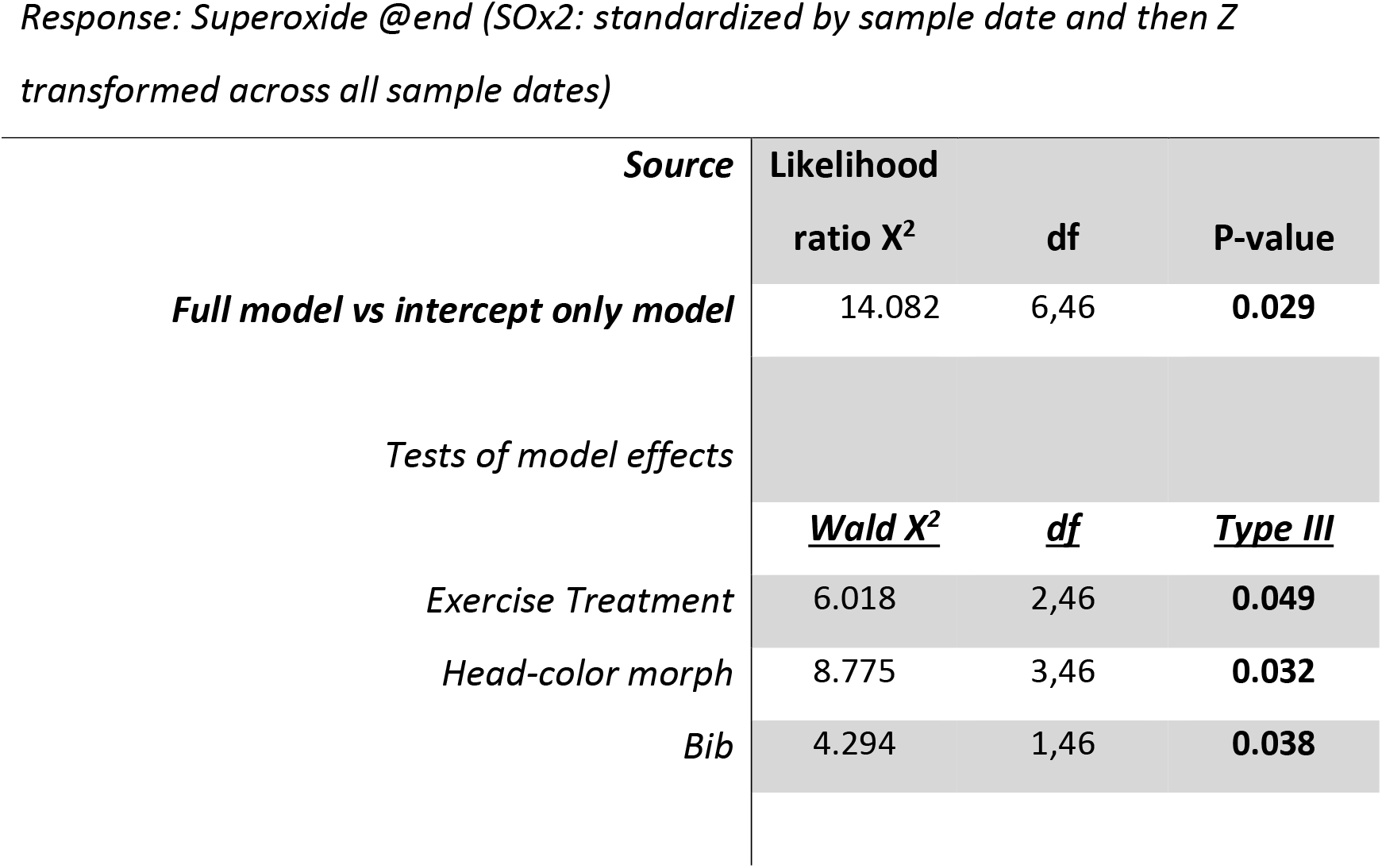
Superoxide. Results of GLM (normal, identity link). Backward elimination of interaction terms between Head-color morph x Exercise (P = 0.263) and Bib x Exercise (P = 0.511) reduced model to main effects only. ∆AICC of reduced (AICC: 153.24) versus full model (AICC: 176.76) = - 23.41. Bold text in p-value column indicates significance at P ≤ 0.05.

**Figure 4.**
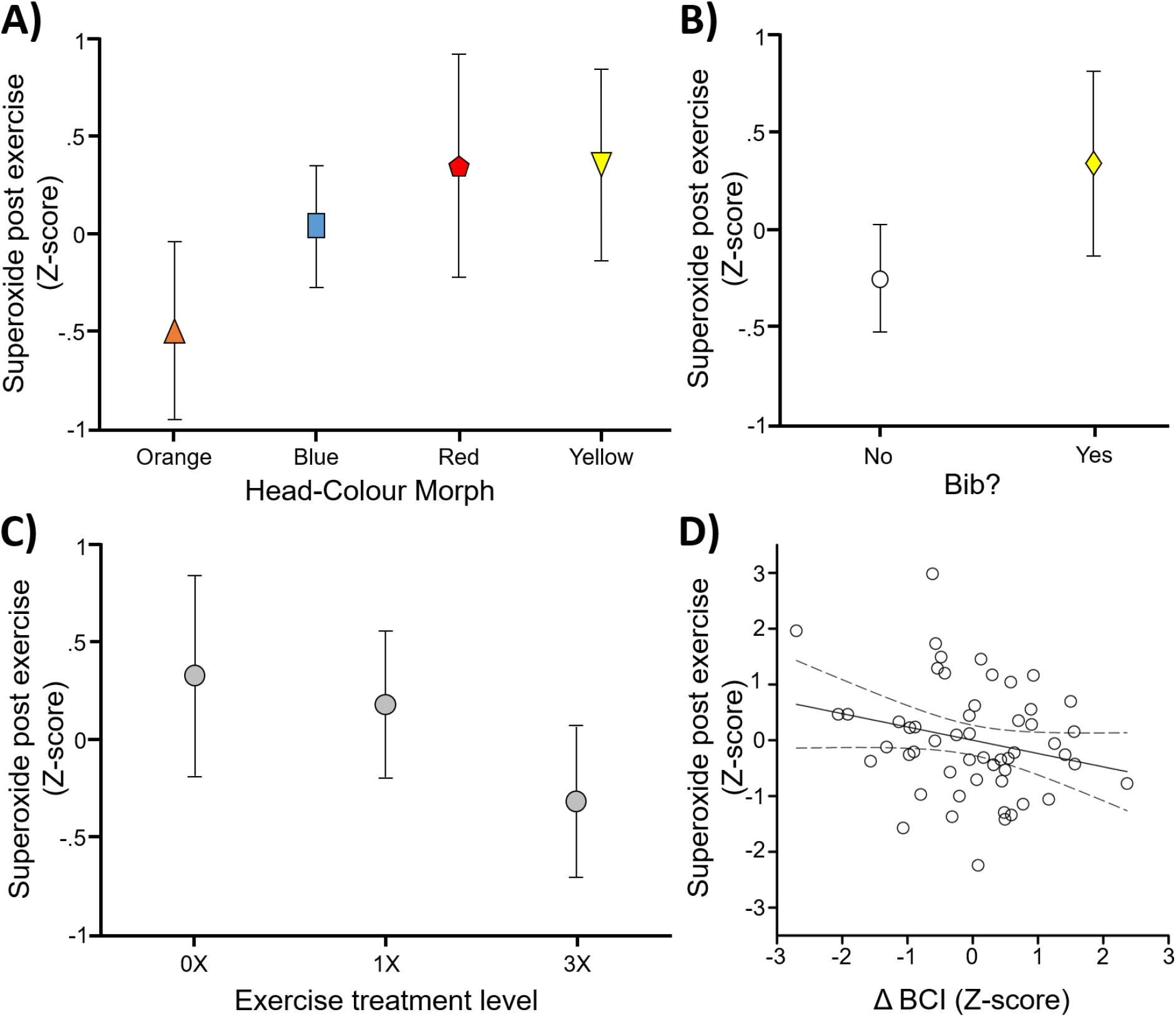
The relationship of superoxide (z-transformed) at the end of the exercise period and A) head-color morphs, B) bib-morphs, C) exercise treatment, and D) change in body condition (z-transformed ∆BCI) (r =−239, F_1,51_ = 3.077, P = 0.085). All symbols are centered on the estimated mean, and the bars represent 95% confidence estimates of the mean from generalized linear models.

##### ROS

ROS was weakly consistent between an individual’s measurement before and after exercise treatment (Spearman’s ρ = 0.263, P = 0.058, ICC = 0.299, 0.033-0.525 95%CI, P = 0.014).

The ROS after exercise treatment and ∆ROS were not affected by exercise treatment, and neither variable was related to initial or post-treatment BCI, ∆BCI, rTL, ∆rTL, head-color, bib-morph, or their interactions with exercise (all P > 0.172). Although not significant at α < 0.05, there was weak evidence of a *positive* relationship between food consumption and ROS at the end of the exercise treatment (Spearman’s ρ = 0.255, P = 0.065) and food consumption and ∆ROS (Spearman’s ρ = 0.263, P = 0.058). There was weak evidence of a *negative* relationship between other ROS *before* exercise treatments began and ∆SVL (Spearman’s ρ = -0.244, P = 0.079), but there was no evidence for a relationship of other ROS with ∆SVL afterwards (Spearman’s ρ = 0.003, P = 0.975).

## Discussion

This experiment, which was inspired by differential condition loss of bibbed males in field studies (Healey and Olsson, 2009; Olsson et al., 2009a), was designed to test the effect of physical exercise and the resultant energy/resource allocation trade-offs on morph-specific telomere dynamics. We manipulated activity levels through enforced exercise treatments to test for morph-specific responses in telomere dynamics in painted dragons (*Ctenophorus pictus*). This experimental approach revealed morph-specific, activity-related lengthening and erosion of telomeres. Telomere dynamics were related to body condition and growth, the latter of which was positively associated with food intake. The differences in telomere dynamics were most evident between bib-morphs. The telomere dynamics of bibbed and non-bibbed males suggests the potential for further longevity trade-offs through the breeding season in this species. The interaction of head color morph with exercise treatment was primarily driven by blue males increasing telomere length with no exercise and shortening when exercised at any level, while the other morphs mostly maintained or lost telomere length. Blue morphs only appeared relatively recently in the sampled population, so we know little about their biology or mating tactics (Olsson et al., 2007b), which limits our scope for interpretation of these results. We do know from previous work that bibbed males and blue males have similarly short telomeres (Rollings et al., 2017) and lower endurance than the other morphs (Tobler et al., 2012), so it is possible that patterns in bibbed males may be relevant for future work on blue males. We, therefore, restrict our discussion to the broader relationships among variables and the differences among bib-morphs because their mating tactics are known and the analyses of bib vs non-bib have higher power than those of the four head-color morphs.

Endurance and physiological performance in lizards is likely to be a polygenic trait, as in humans allelic variation at over 20 loci contributes to endurance performance phenotype (Williams and Folland, 2008) and the ability to respond to training (Mann et al., 2014). It seems likely such allelic variation is differentially aggregated in bibbed and non-bibbed morphs. Differential energy strategies are likely to underpin different reproductive tactics of all morphs. However, in the lab, animals usually do not have the same opportunity for activity as in the wild, where they must forage for food, escape predators, and find and defend territories. In our no-exercise (0x) treatment, bibbed males maintained or increased BCI, telomere length, and body length simultaneously. With increasing energetic demands of exercise, bibbed males exhibited reduced growth, condition and lost telomere length.

When there are multiple solutions to balance the costs and benefits of various mating/fertilization strategies, alternative phenotypes and polymorphisms are likely to evolve (Neff and Svensson, 2013; Smith, 1982; West-Eberhard, 1983). These alternative reproductive tactics (ARTs) are often associated with physiological phenotypes and energy expenditure strategies (Miles et al., 2007; Soulsbury, 2019), growth rates (Hazard et al., 2019), and color-polymorphisms (Olsson et al., 2013; Wellenreuther et al., 2014) that reflect a balance between pre- and postcopulatory traits (Parker et al., 2013).

Both bibbed and red-headed males appear to invest more in traits that aid in precopulatory sexual selection. Red and bibbed males end to be more aggressive, winning male-male contests (Healey et al., 2007; McDiarmid et al., 2017). Red males have higher blood testosterone concentrations and ROS concentrations (Olsson et al., 2007a; Olsson et al., 2009c), and bibbed males have higher metabolic rates and lower endogenous antioxidant activity and endurance than other morphs (Friesen et al., 2019; Friesen et al., 2017a; Tobler et al., 2012). Yellow and non-bibbed males seem to have similar reproductive tactics that include having larger testes (Olsson et al., 2009b), and inseminating more and faster sperm (Friesen et al., 2020) but with significantly shorter copulation durations than either red or bibbed males (Friesen et al., 2020; Olsson et al., 2009b). Thus, red and bibbed males are similar to each other, as are yellow and non-bibbed males. Yellow males have 3x greater paternity success in head-to-head sperm competition trials over red males (Olsson et al., 2009b). However, field studies indicate that bibbed males do not lose paternity to neighboring males, which is probably because of effective territorial defense and mate guarding (Healey and Olsson, 2009; Olsson et al., 2009a). Bibbed males quickly lose body condition in the field, which may indicate high activity levels due to territorial defense and/or lower foraging time (Healey and Olsson, 2009; Olsson et al., 2009a).

In line with our predictions and the previous work described above, the aggressive bibbed phenotype seems to pay the cost of telomere attrition when they sustain exhaustive, high endurance activities. Males with bibs gained telomere length with no exercise (0X) and lost telomere length with increased exercise (3X), while those without the bib showed the opposite pattern. Across all males, body condition and telomere dynamics were correlated (**Fig 2**). Positive changes in body condition across exercise treatments and morphs were qualitatively associated with increases in telomere length, a trend especially apparent in bib-morphs (**Fig 1A and B**). The behavior of bibbed males matches their territorial role, as they are the first to be aggressive during territorial-contests and have faster reaction times than non-bibbed males (McDiarmid et al., 2017; Tobler et al., 2012). Non-bibbed males have better endurance, tend to copulate for shorter periods, but inseminate more andfaster sperm than bibbed males (Friesen et al., 2020), suggesting that non-bibbed males employ an alternative, sneaker/satellite male strategy rather than monopolizing females (sensu Smith, 1982; Smith and Price, 1973). It is difficult to say whether the exercise treatment we employed better represents the energy expenditure of a territorial or a sneaker strategy. Nevertheless, the exercise levels do represent incremental increases in energy expenditure, which may force and expose allocation trade-offs. The association of BCI and telomere loss across exercise treatments within bib-morphs is congruent with patterns of greater-than-average mass loss of bibbed males in the wild (Healey and Olsson, 2009; Olsson et al., 2009a) where territorial defense gives bibbed males a paternity advantage (Olsson et al., 2009a), despite their smaller ejaculates. Displaying a bib seems to incur a “social cost” from more frequent challenges to and by rivals and higher energetic expenditure, resulting in poorer condition (Olsson et al., 2007b). Bibbed males have higher resting metabolic rates than non-bibbed males (Friesen et al., 2017a), lower endogenous antioxidant protection (Friesen et al., 2019), and higher superoxide levels (this study), which along with decreasing body condition may also contribute to telomere loss in high-activity bibbed males. These various “social” and physiological costs may, in part, explain decreases in the frequency of bibbed morphs across some years (Healey and Olsson, 2009).

Among lizards, body size and aggressiveness are generally positively associated with contest success (Olsson and Madsen, 1998), but growth rate and aggression are often negatively correlated with survival (Olsson and Shine, 2002; Stamps, 2007). In this study, across all males (regardless of treatment), growth correlated with shorter final telomeres, while those males that shrunk in size or did not grow tended to maintain or even increase telomere length—the bibbed males invested in higher growth rates overall, which is congruent with their aggressive tactics. Bibbed males maintained body condition while they also maximized growth rate with no-exercise (0X), which is probably a lower level of activity and energy expenditure than free-living lizards. Non-bibbed males represented the majority of the males that did not grow. The growth rate and final telomere length are likely interrelated and probably reflect morph-specific strategies on a fast-slow pace of life continuum.

Rates of longevity and senescence can differ among morphs (Zamudio and Sinervo, 2000). Color-polymorphic species are a valuable tool in resolving the interplay between sexually selected and life-history traits because the polymorphism is a convenient visual-code for the associated behaviors and physiology in an otherwise common genetic background (Stuart-Fox et al., 2020). Longevity is predicted to be influenced by oxidative stress (Kirkwood, 2017); however, although we know that annual mortality is 90% variation of within-season longevity has not been investigated in the field for this species. We found no direct link between either SOx or other ROS and telomere erosion (highest correlation coefficient |0.192|, all P < 0.168), but males that grew more tended to have higher superoxide and other ROS levels. As a group, bibbed males grew significantly more and had higher superoxide— suggesting indirect links between telomere length and growth, energy utilization patterns, and ROS levels.

These results imply different energy pathways for growth and bodily maintenance, possibly extending to telomerase expression and activity and antioxidant protection (which we did not measure). We predicted a negative relationship between growth and telomere dynamics. Approximately 36% of the males decreased in length or did not grow. Growth was negatively related to telomere length at both the beginning and end of the experiment but not to ∆rTL, suggesting that an individual’s prior investments in maintenance, early in the breeding season or earlier in life, set up growth and telomere length trajectories (e.g., Nettle et al., 2015; Parolini et al., 2015). Growth was positively related to food consumption, suggesting that energy and critical resources (e.g., calcium or vitamins dusted on mealworms) limit growth. However, food consumption was not related to increased body condition (i.e., residual body mass that is independent of body length) or telomere dynamics. Indeed, across all the males, ignoring exercise treatments, the relationships between telomere length was negatively correlated with growth but positively correlated with improved body condition. This result may, in part, be explained by the weak evidence of a negative correlation between BCI and superoxide (r = -0.239, *one-tailed* P = 0.043), suggesting a mechanistic link between BCI and telomere protection from oxidative damage (Barnes et al., 2019; von Zglinicki, 2002).

In this species, Friesen et al. (2019) demonstrated a significant, indirect negative link between superoxide and body condition by describing a negative relationship between superoxide levels and superoxide dismutase (SOD the enzyme that quenches SOx) activity, and a positive relationship of SOD with BCI. These relationships were also bib-morph specific, with bibbed males showing lower SOD activity than non-bibbed males, and sperm that were more sensitive to superoxide (Friesen et al., 2019). Interestingly, in this study, we also found tentative evidence that food consumption may be positively related to ROS, which, if true, might be due to the high-fat content of mealworms (~ 32% fat (Ravzanaadii et al., 2012)) providing more substrate for lipid peroxidation and concomitant free-radical cascade (Halliwell and Gutteridge, 2015). However, others have found that simply eating and digesting food comes at an oxidative cost (Butler et al., 2016). Mitochondrial function and ROS production are complex and depend on substrates (Castro et al., 2015; Martos-Sitcha et al., 2017) that may differentially affect the fitness of different mitochondrial haplotypes (Aw et al., 2018; Ballard and Youngson, 2015). In ectotherms, shifts from aerobic to anaerobic metabolism may also reduce ROS production in mitochondria, relieving oxidative stress (Pérez-Jiménez et al., 2012). We found no evidence of an interaction between food consumption and bib-morph on ROS (all P > 0.187), but the link between energy use, diet, and oxidative stress may have important implications for life-history and reproductive strategies.

The relationships between growth, condition, and telomere length are reversed in non-bibbed males and may be linked to different reproductive tactics and energy allocation. The increased exercise led to decreased growth but increased telomere length in non-bibbed males. In some lizards, sprint performance aids in territorial defense and, thus, paternity (e.g., Garland Jr et al., 1990; Husak et al., 2006; Husak et al., 2008; Robson and Miles, 2000). While sprint performance and endurance are positively correlated in some lizard species, others exhibit a trade-off between these traits (e.g., Vanhooydonck et al., 2001). Given the circumstantial evidence that non-bibbed males engage in sneaker-male tactics, their regular movement among territories and fleeing from territorial males may generate selection on their endurance performance. Furthermore, male size is important in territorial defense but less so in sneaker males, which might even benefit from a reduction in size and the consequently lower total metabolic costs (Blanckenhorn, 2000). However, in side-blotched lizards (*Uta stansburiana*), larger orange territorial males have higher endurance than smaller yellow sneakers (Hazard et al., 2019). In painted dragons, higher endurance of non-bibbed morphs and maintenance of longer telomeres is suggestive of increased survival and, thus, a longer reproductive life (Healey and Olsson, 2009; Olsson et al., 2009a). The higher metabolism, aggression, and growth rate of bibbed males are likely to be associated with reproductive success early in the breeding season (Healey and Olsson, 2008), which may select for sperm longevity through posthumous paternity (Olsson et al., 2009b; Zamudio and Sinervo, 2000). Indeed, bibbed males have longer sperm telomeres (Friesen et al., 2020), which may be an advantage in sperm longevity and, thus, female sperm storage. However, the implications of sperm telomere length for male fitness have only recently been investigated. The potential for sperm telomere length to influence sperm competitiveness or biases in female sperm storage are exciting ideas remaining to be tested (Friesen et al., 2020; Olsson et al., 2018b; Olsson et al., 2017). These studies could extend to potential transgenerational links to offspring fitness, which could be explained by sperm telomeres acting as biomarkers of, for example, DNA fragmentation or other free radical-induced sperm damage sublethally affecting development and offspring performance (Pauliny et al., 2018). This work highlights the utility of combining complementary, controlled lab-based experimental work with fieldwork to generate and iteratively test hypotheses in the wild.

## Authors’ Contributions

All authors contributed critically to the drafts and gave final approval for publication. CRF and MO conceived the ideas; CRF, MO, JS, and MG designed experiments; CRF, MW, NR, JS, MG, and MO collected the samples; CRF, MW, NR, CMW, and MO analyzed the data; CRF, CMW, and MO led the writing of the manuscript.

## Acknowledgements

We thank the Australian Research Council (MO), the National Science Foundation (CRF), and the University of Wollongong (CRF) for financial support, E Snaith for logistic field support, G Ljungström for inspiring this project, C Bezzina for help with husbandry and as lizard-swim-coach, and EJ Uhrig for sagacious editorial comments.

## Data Accessibility

Data will be made available in the Dryad Digital Repository upon acceptance of the manuscript.

## Supplemental tables Exercise exp dragons

**Table 1:** ∆ rTL. Results of GLM (normal, identity link function) and parameter estimates. Bold text in p‐value column indicates significance at P ≤ 0.05.

| Source | Likelihood |  |  |
| --- | --- | --- | --- |
| | ratio $X^2$ | df | P-value |
| **Full model vs intercept only model** | 25.356 | 14,38 | **0.031** |
| **Tests of model effects** |  |  |  |
|  | **<u>Wald $X^2$</u>** | **<u>df</u>** | **<u>Type III</u>** |
| Exercise Treatment | 1.341 | 2,38 | .512 |
| Head-color morph | 5.408 | 3,38 | .144 |
| Bib | .059 | 1,38 | .808 |
| Exerc.Treat. * HC-Morph | 47.532 | 6,38 | **<0.001** |
| Exerc.Treat. * Bib | 19.330 | 2,38 | **<0.001** |

Parameter estimates for $\Delta$ Telomeres (Z-transformed (rTL (End-Beginning)))
| Parameter | B | Std. Error | 95% Wald Confidence Interval |  | Hypothesis Test |  |  |
| --- | --- | --- | --- | --- | --- | --- | --- |
|  |  |  | Lower | Upper | Wald Chi-Square | df | Sig. |
| (Intercept) | -0.076 | 0.4143 | -0.888 | 0.736 | 0.034 | 1 | 0.854 |
| [Exercisetreat=0X] | 0.050 | 0.4394 | -0.811 | 0.911 | 0.013 | 1 | 0.910 |
| [Exercisetreat=1X] | -0.199 | 0.5361 | -1.250 | 0.851 | 0.138 | 1 | 0.710 |
| [Exercisetreat=3X] | 0<sup>a</sup> |  |  |  |  |  |  |
| [MORPHS=BLUE ] | -0.775 | 0.7429 | -2.231 | 0.681 | 1.087 | 1 | 0.297 |
| [MORPHS=ORANGE] | -1.314 | 0.5749 | -2.441 | -0.187 | 5.225 | 1 | **0.022** |
| [MORPHS=RED ] | -1.002 | 0.5334 | -2.047 | 0.043 | 3.529 | 1 | 0.060 |
| [MORPHS=YELLOW] | 0<sup>a</sup> |  |  |  |  |  |  |
| [BIB=NO ] | 1.384 | 0.5107 | 0.383 | 2.385 | 7.346 | 1 | **0.007** |
| [BIB=YES] | 0<sup>a</sup> |  |  |  |  |  |  |
| [Exercisetreat=0X] * | 2.027 | 0.8195 | 0.420 | 3.633 | 6.116 | 1 | **0.013** |
| [MORPHS=BLUE ] |  |  |  |  |  |  |  |
| [Exercisetreat=0X] * | 0.649 | 0.6845 | -0.693 | 1.990 | 0.898 | 1 | 0.343 |
| [MORPHS=ORANGE] |  |  |  |  |  |  |  |
| [Exercisetreat=0X] * | 1.663 | 0.6225 | 0.443 | 2.883 | 7.134 | 1 | **0.008** |
| [MORPHS=RED ] |  |  |  |  |  |  |  |
| [Exercisetreat=0X] * | 0<sup>a</sup> |  |  |  |  |  |  |
| [MORPHS=YELLOW] |  |  |  |  |  |  |  |
| [Exercisetreat=1X] * | 0.550 | 0.7990 | -1.016 | 2.116 | 0.474 | 1 | 0.491 |
| [MORPHS=BLUE ] |  |  |  |  |  |  |  |
| [Exercisetreat=1X] * | 1.971 | 0.6656 | 0.667 | 3.276 | 8.771 | 1 | **0.003** |
| [MORPHS=ORANGE] |  |  |  |  |  |  |  |
| [Exercisetreat=1X] * | 1.822 | 0.7645 | 0.324 | 3.321 | 5.681 | 1 | **0.017** |
| [MORPHS=RED ] |  |  |  |  |  |  |  |
| [Exercisetreat=1X] * | 0<sup>a</sup> |  |  |  |  |  |  |
| [MORPHS=YELLOW] |  |  |  |  |  |  |  |
| [Exercisetreat=3X] * | 0<sup>a</sup> |  |  |  |  |  |  |
| [MORPHS=BLUE ] |  |  |  |  |  |  |  |
| [Exercisetreat=3X] * | 0<sup>a</sup> |  |  |  |  |  |  |
| [MORPHS=ORANGE] |  |  |  |  |  |  |  |
| [Exercisetreat=3X] * | 0<sup>a</sup> |  |  |  |  |  |  |
| [MORPHS=RED ] |  |  |  |  |  |  |  |
| [Exercisetreat=3X] * | 0<sup>a</sup> |  |  |  |  |  |  |
| [MORPHS=YELLOW] |  |  |  |  |  |  |  |
| [Exercisetreat=0X] * | -2.484 | 0.5822 | -3.625 | -1.342 | 18.197 | 1 | **0.000** |
| [BIB=NO ] |  |  |  |  |  |  |  |
| [Exercisetreat=0X] * | 0<sup>a</sup> |  |  |  |  |  |  |
| [BIB=YES] |  |  |  |  |  |  |  |
| [Exercisetreat=1X] * | -1.511 | 0.5845 | -2.657 | -0.366 | 6.686 | 1 | **0.010** |
| [BIB=NO ] |  |  |  |  |  |  |  |
| *[Exercisetreat=1X] \**<br>*[BIB=YES]* | 0<sup>a</sup> |  |  |  |  |  |  |
| *[Exercisetreat=3X] \**<br>*[BIB=NO ]* | 0<sup>a</sup> |  |  |  |  |  |  |
| *[Exercisetreat=3X] \**<br>*[BIB=YES]* | 0<sup>a</sup> |  |  |  |  |  |  |
| *(Scale)* | .608<sup>b</sup> | 0.1181 | 0.416 | 0.890 |  |  |  |

*Model: (Intercept), Exercisetreat, MORPHS, BIB, Exercisetreat \* MORPHS, Exercisetreat \* BIB*
*a. Set to zero because this parameter is redundant.*
*b. Maximum likelihood estimate.*

**Table 2:** ∆ BCI. Results of GLM (normal, identity link function) and parameter estimates. Bold text in the p‐value column indicates significance at P ≤ 0.05.

| *Source* | **Likelihood** |  |  |
| --- | --- | --- | --- |
|  | **ratio X<sup>2</sup>** | **df** | **P-value** |
| ***Full model vs intercept only model*** | 36.906 | 14,38 | **0.001** |
| *Tests of model effects* |  |  |  |
|  | ***Wald X<sup>2</sup>*** | ***df*** | ***Type III*** |
| *Exercise Treatment* | 2.777 | 2,38 | 0.250 |
| *Head-color morph* | 4.612 | 3,38 | 0.203 |
| *Bib* | 2.060 | 1,38 | 0.151 |
| *Exerc.Treat. \* HC-Morph* | 21.402 | 6,38 | **0.002** |
| *Exerc.Treat. \* Bib* | 18.777 | 2,38 | **<0.001** |

***Parameter estimates for $\Delta$ BCI (Z-transformed (BCI (End-Beginning)))***
| *Parameter* | **B** | **Std. Error** | **95% Wald**<br>**Confidence**<br>**Interval** | **Hypothesis Test** |
| --- | --- | --- | --- | --- |

|  |  |  | Lower | Upper | Wald<br>Chi-Square | df | Sig. |
| --- | --- | --- | --- | --- | --- | --- | --- |
| *(Intercept)* | -0.730 | 0.2794 | -1.277 | -0.182 | 6.815 | 1 | **0.009** |
| *[Exercisetreat=0X]* | 0.550 | 0.3535 | -0.142 | 1.243 | 2.424 | 1 | 0.119 |
| *[Exercisetreat=1X]* | -0.020 | 0.7774 | -1.543 | 1.504 | 0.001 | 1 | 0.980 |
| *[Exercisetreat=3X]* | 0a |  |  |  |  |  |  |
| *[MORPHS=BLUE ]* | 0.638 | 0.5021 | -0.346 | 1.622 | 1.615 | 1 | 0.204 |
| *[MORPHS=ORANGE]* | 0.493 | 0.5507 | -0.586 | 1.573 | 0.802 | 1 | 0.370 |
| *[MORPHS=RED ]* | 0.353 | 0.5777 | -0.779 | 1.486 | 0.374 | 1 | 0.541 |
| *[MORPHS=YELLOW]* | 0a |  |  |  |  |  |  |
| *[BIB=NO ]* | 1.088 | 0.4127 | 0.280 | 1.897 | 6.956 | 1 | **0.008** |
| *[BIB=YES]* | 0a |  |  |  |  |  |  |
| *[Exercisetreat=0X] \** | 0.724 | 0.5899 | -0.432 | 1.880 | 1.507 | 1 | 0.220 |
| *[MORPHS=BLUE ]* |  |  |  |  |  |  |  |
| *[Exercisetreat=0X] \** | 0.351 | 0.6003 | -0.826 | 1.528 | 0.342 | 1 | 0.559 |
| *[MORPHS=ORANGE]* |  |  |  |  |  |  |  |
| *[Exercisetreat=0X] \** | -0.592 | 0.6568 | -1.879 | 0.696 | 0.812 | 1 | 0.368 |
| *[MORPHS=RED ]* |  |  |  |  |  |  |  |
| *[Exercisetreat=0X] \** | 0a |  |  |  |  |  |  |
| *[MORPHS=YELLOW]* |  |  |  |  |  |  |  |
| *[Exercisetreat=1X] \** | -0.824 | 0.7933 | -2.379 | 0.731 | 1.079 | 1 | 0.299 |
| *[MORPHS=BLUE ]* |  |  |  |  |  |  |  |
| *[Exercisetreat=1X] \** | -0.648 | 0.7814 | -2.180 | 0.883 | 0.688 | 1 | 0.407 |
| *[MORPHS=ORANGE]* |  |  |  |  |  |  |  |
| *[Exercisetreat=1X] \** | 0.375 | 0.7409 | -1.077 | 1.827 | 0.256 | 1 | 0.613 |
| *[MORPHS=RED ]* |  |  |  |  |  |  |  |
| *[Exercisetreat=1X] \** | 0a |  |  |  |  |  |  |
| *[MORPHS=YELLOW]* |  |  |  |  |  |  |  |
| *[Exercisetreat=3X] \** | 0a |  |  |  |  |  |  |
| *[MORPHS=BLUE ]* |  |  |  |  |  |  |  |
| *[Exercisetreat=3X] \** | 0a |  |  |  |  |  |  |
| *[MORPHS=ORANGE]* |  |  |  |  |  |  |  |
| *[Exercisetreat=3X] \** | 0a |  |  |  |  |  |  |
| *[MORPHS=RED ]* |  |  |  |  |  |  |  |
| *[Exercisetreat=3X] \** | 0a |  |  |  |  |  |  |
| *[MORPHS=YELLOW]* |  |  |  |  |  |  |  |
| *[Exercisetreat=0X] \** | -1.821 | 0.4621 | -2.727 | -0.915 | 15.530 | 1 | **0.000** |
| *[BIB=NO ]* |  |  |  |  |  |  |  |
| *[Exercisetreat=0X] \** | 0a |  |  |  |  |  |  |
| *[BIB=YES]* |  |  |  |  |  |  |  |
| *[Exercisetreat=1X] \** | -0.426 | 0.6786 | -1.756 | 0.904 | 0.394 | 1 | 0.530 |
| *[BIB=NO ]* |  |  |  |  |  |  |  |
| *[Exercisetreat=1X] \** | 0a |  |  |  |  |  |  |
| *[BIB=YES]* |  |  |  |  |  |  |  |
| *[Exercisetreat=3X] \** | 0a |  |  |  |  |  |  |
| *[BIB=NO ]* |  |  |  |  |  |  |  |
| *[Exercisetreat=3X] \** | 0a |  |  |  |  |  |  |
| *[BIB=YES]* |  |  |  |  |  |  |  |
| *(Scale)* | .489b | 0.0950 | 0.334 | 0.716 |  |  |  |

Model: (Intercept), Exercisetreat, MORPHS, BIB, Exercisetreat \* MORPHS, Exercisetreat \* BIB
a. Set to zero because this parameter is redundant.
b. Maximum likelihood estimate.

**Table 3:**
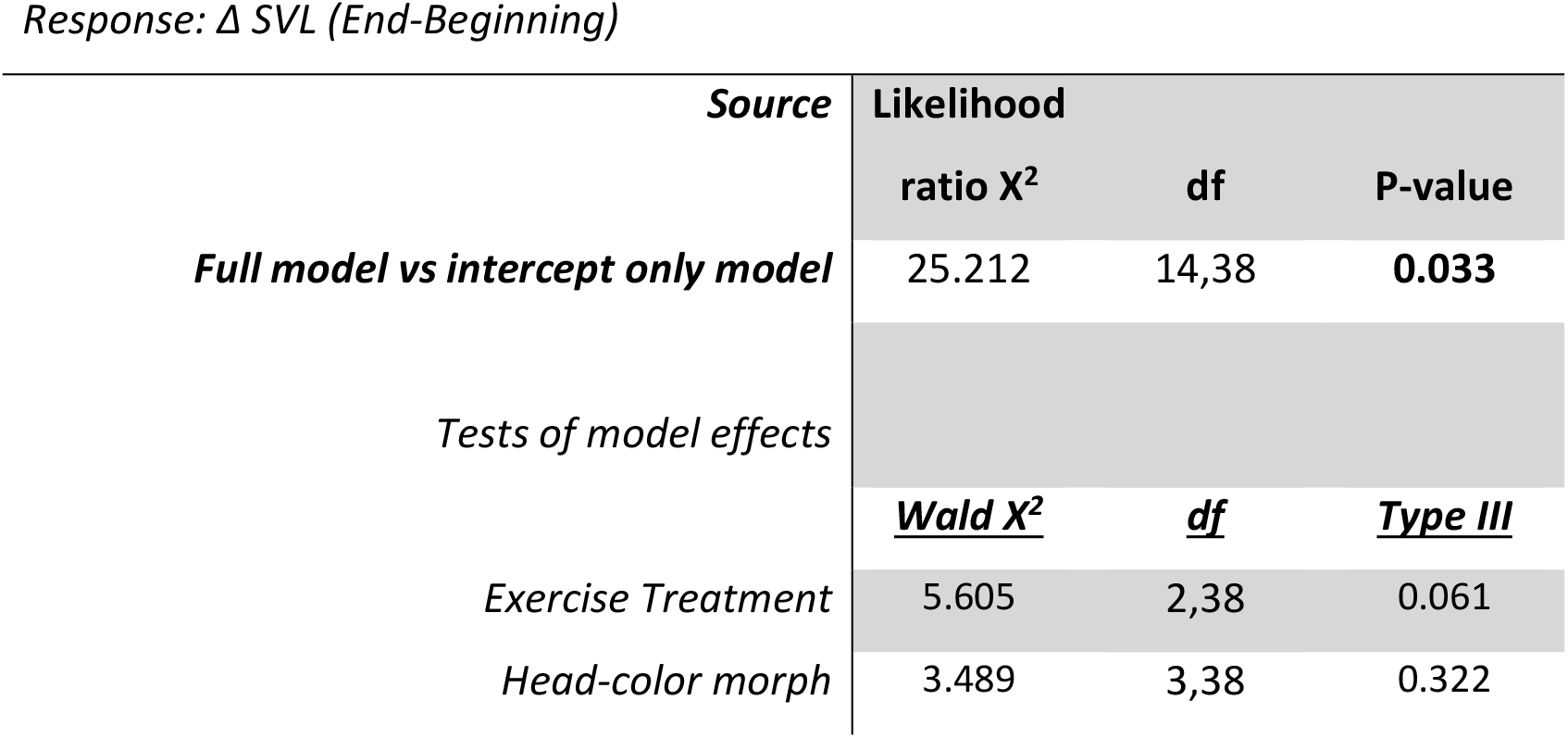
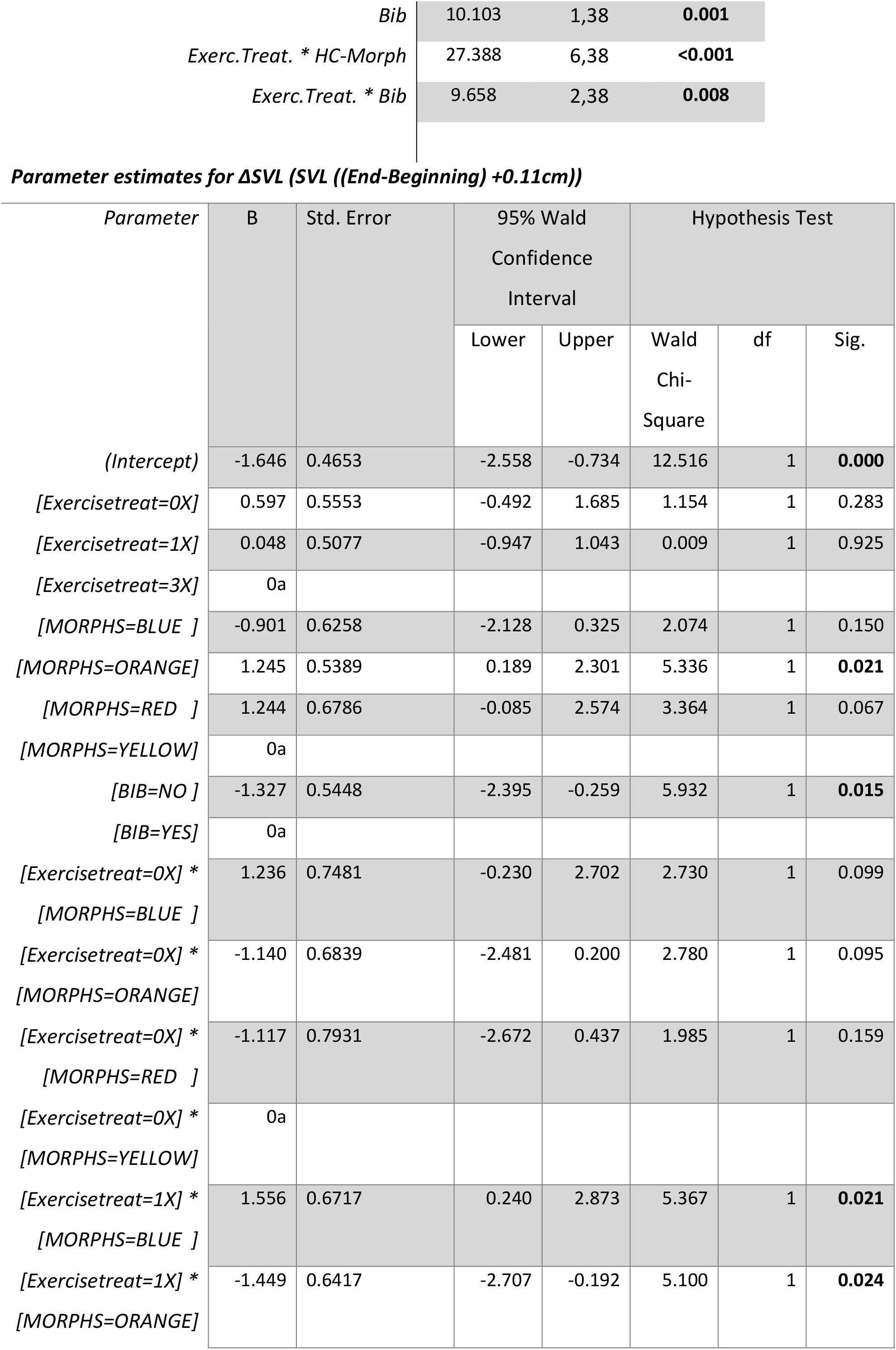
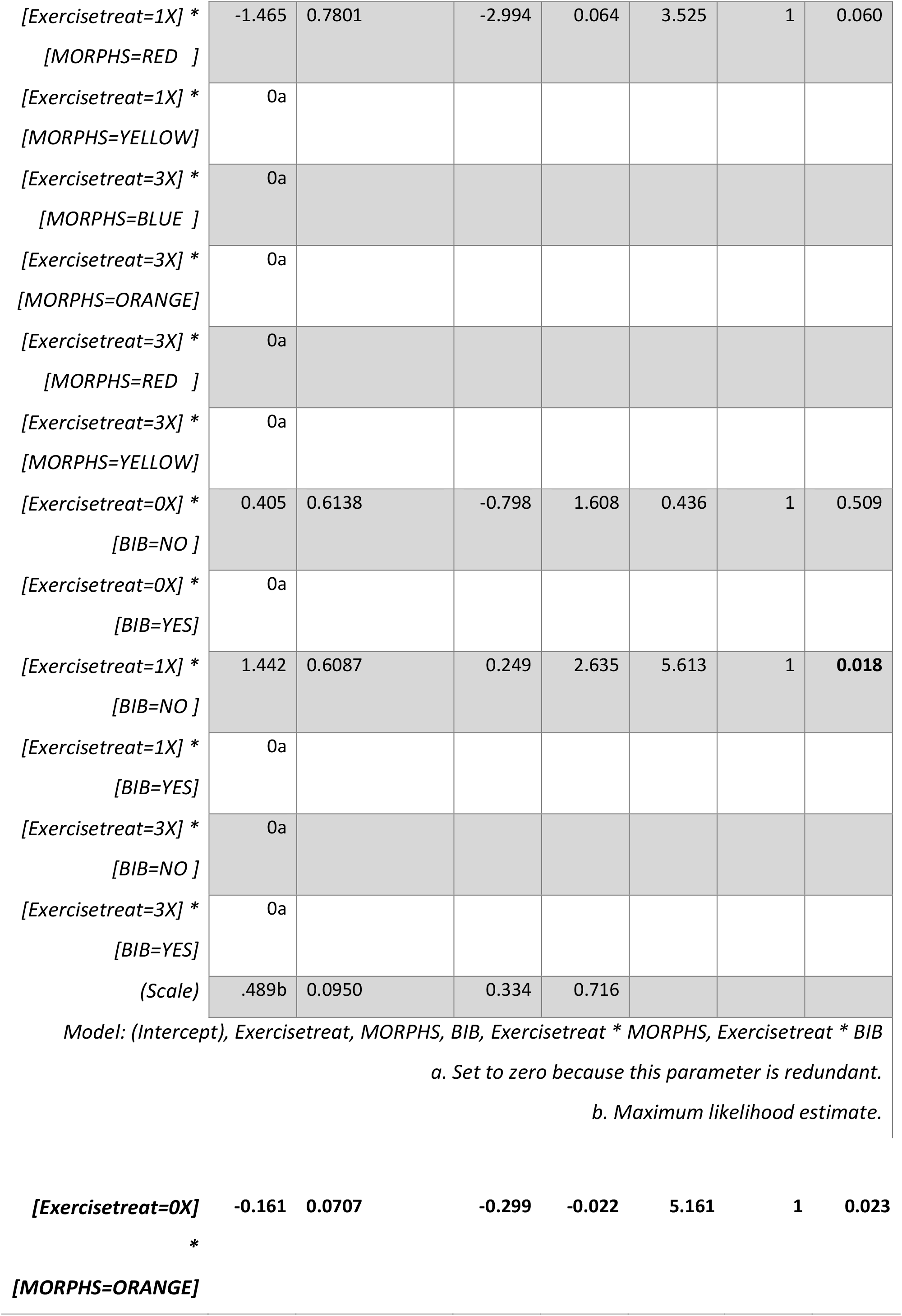
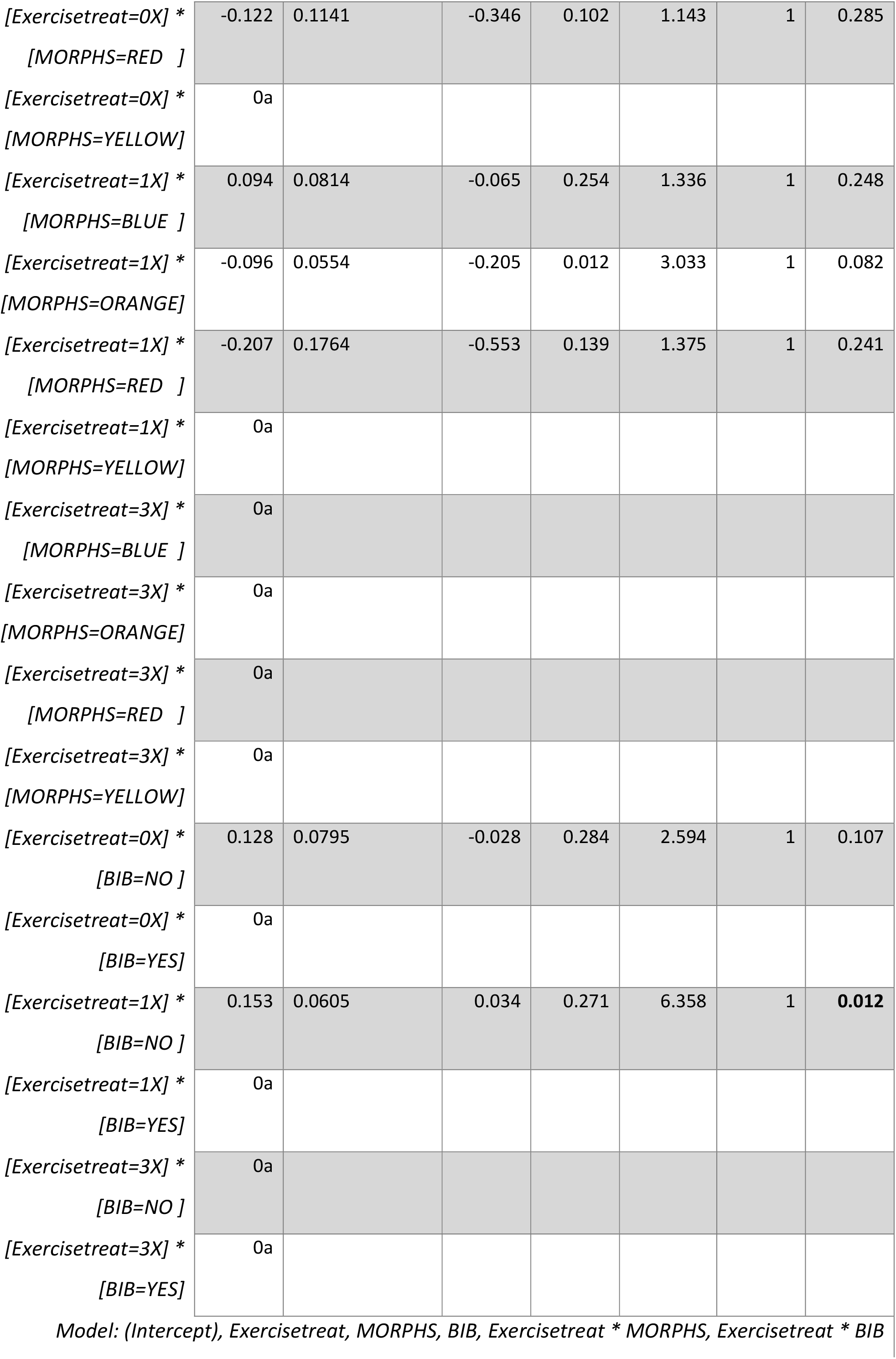
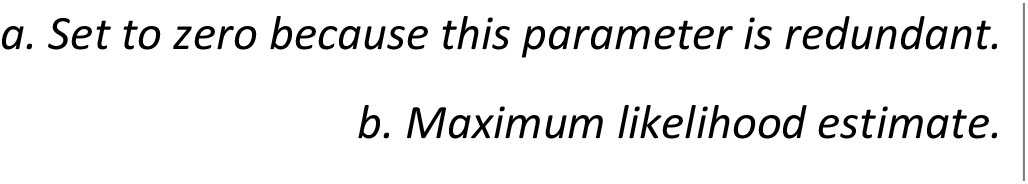
∆ SVL. Results of GLM (gamma, log link function) and parameter estimates. Deviance value = 36.639, degrees of freedom = 38, Deviance/df = 0.964, indicating that the model is not over‐dispersed. Bold text in the p‐value column indicates significance at P ≤ 0.05.

**Table 4:** Superoxide. Results of GLM (normal, identity link) and parameter estimates. Backward elimination of interaction terms between Head‐color morph x Exercise (P = 0.263) and Bib x Exercise (P = 0.511) reduced model to main effects only. ∆AICC of reduced (AICC: 153.24) versus full model (AICC: 176.76) = ‐23.41. Bold text in p‐value column indicates significance at P ≤ 0.05.

| Source | Likelihood |  |  |
| --- | --- | --- | --- |
| | ratio $X^2$ | df | P-value |
| **Full model vs intercept only model** | 14.082 | 6,46 | **0.029** |
| **Tests of model effects** |  |  |  |
|  | **<u>Wald $X^2$</u>** | **<u>df</u>** | **<u>Type III</u>** |
| Exercise Treatment | 6.018 | 2,46 | **0.049** |
| Head-color morph | 8.775 | 3,46 | **0.032** |
| Bib | 4.294 | 1,46 | **0.038** |

Parameter estimates for Mitochondrial superoxide @end (Z transformed after standardized by date)
| Parameter | B | Std. Error | 95% Wald Confidence Interval |  | Hypothesis Test |  |  |
| --- | --- | --- | --- | --- | --- | --- | --- |
|  |  |  | Lower | Upper | Wald Chi-Square | df | Sig. |
| (Intercept) | 0.264 | 0.3573 | -0.437 | 0.964 | 0.544 | 1 | 0.461 |
| [Exercisetreat=0X] | 0.642 | 0.3107 | 0.034 | 1.251 | 4.276 | 1 | **0.039** |
| [Exercisetreat=1X] | 0.495 | 0.2480 | 0.009 | 0.981 | 3.982 | 1 | **0.046** |
| [Exercisetreat=3X] | 0a |  |  |  |  |  |  |
| [MORPHS=BLUE ] | -0.315 | 0.2774 | -0.859 | 0.228 | 1.292 | 1 | 0.256 |
| *[MORPHS=ORANGE]* | -0.849 | 0.3315 | -1.499 | -0.199 | 6.561 | 1 | **0.010** |
| *[MORPHS=RED ]* | -0.005 | 0.3343 | -0.660 | 0.651 | 0.000 | 1 | 0.989 |
| *[MORPHS=YELLOW]* | 0a |  |  |  |  |  |  |
| *[BIB=NO ]* | -0.586 | 0.2828 | -1.140 | -0.032 | 4.294 | 1 | **0.038** |
| *[BIB=YES]* | 0a |  |  |  |  |  |  |
| *(Scale)* | .752b | 0.1461 | 0.514 | 1.101 |  |  |  |

*Model: (Intercept), Exercisetreat, MORPHS, BIB*
*a. Set to zero because this parameter is redundant.*
*b. Maximum likelihood estimate.*

